# A novel microcosm for recruiting inherently competitive biofertilizer-candidate microorganisms from soil environments

**DOI:** 10.1101/2020.09.03.274811

**Authors:** S Pittroff, S Olsson, Ashlea Doolette, R. Greiner, A.E. Richardson, M Nicolaisen

## Abstract

Fertilizer phosphorus (P) is both a necessary crop nutrient and finite resource, necessitating the development of innovative solutions for P fertilizer efficiency and recycling in agricultural systems. *Myo*-inositol hexa*kis*phosphate (phytate) and its lower order derivatives constitute the majority of identified organic P in many soil types and has been shown to accumulate with increasing application of P fertilizer. Phytate is only poorly available to plants, and in alkaline soils it often precipitated as even more unavailable calcium (Ca)-phytate. Incorporating phytase-producing biofertilizers (i.e., microbial-based products with capacity to mineralize phytate) into soil presents a viable and environmentally acceptable way of utilizing P from phytate, whilst reducing the need for mineral P application. Here we present an in-soil microcosm that utilizes precipitated Ca-phytate to recruit microorganisms with degradation activity towards phytate *in solum*. Our results show both direct and indirect evidence for Ca-phytate mineralization *in vitro* and *in solum*. Furthermore, the abundance of bacteria recruited was measured via 16S rRNA gene copy number, as was three genes relating to organic P degradation; *phoX* and *phoD* phosphatases and the BPP (β-propeller phytase) gene. Amplicon sequencing as well as BioLog catabolism studies show that microcosm treatments containing the ‘bait’ Ca-phytate, recruited a different set of microorganisms when compared to controls. These Ca-phytate microcosms recruited mainly Actinobacteria, Firmicutes, and Proteobacteria, and the genus *Streptomyces* was specifically enriched. We conclude that our microcosm presents an innovative approach for isolating soil microorganisms with the potential to degrade precipitated phytate *in solum* and represents a new isolation method with the potential to isolate inherently robust biofertilizer candidates directly from target soils.

## 1. Introduction

Biofertilizers are considered to be a cost effective and ecofriendly alternative to chemical fertilizers that can be used in the place of, or in combination with, conventional fertilizers as a means to enhance soil fertility (Singh et al. 2011, Bhardwaj et al. 2014). One obstacle that is consistently encountered is the translation of desirable traits identified from *in-vitro* studies to consistent performance or persistence of those traits in soil (Richardson et al. 2001, Rengel and Marschner 2005). One of the key challenges is the complex and competitive nature of soils, which harbor highly diverse and abundant communities of indigenous microorganisms. This preexisting soil ecosystem represents a barrier for the establishment of specific introduced strains of interest (Malusa et al. 2012, Raynaud and Nunan 2014). While studies based on pure cultures or with a few interacting strains have revealed important insight into the physiology of soil microorganisms, there is an overwhelming need to better understand functional capacities in real soil systems (de Menezes et al. 2017). This limited understanding of soil ecology and function represents a major impediment to the effectiveness of biofertilizers that are expected to perform specific tasks under these complex conditions. Focus on new isolation techniques to obtain *in solum*-compatible microorganisms with plant-growth-promoting capabilities and improved in-field performance has therefore increased (Ghodsalavi et al. 2017, Nicolaisen et al. 2018).

Organic P in soil is a significant pool of P that is currently not considered in modern agricultural production (Cordell et al. 2011, Cordell and White 2011, Stutter et al. 2012, George et al. 2016). The major identifiable form of organic P found in soils is phytate, or *myo*-inositol 1,2,3,4,5,6-hexa*kis*phosphate (IP_6_). It represents a substantial pool of P in soils that is largely not available to plants (Turner et al. 2002, Mullaney 2007, Yan et al. 2014). Phytate reacts readily in soil through adsorption reactions, and has been shown to form precipitates with iron and aluminum (Fe, Al) in acidic soils or calcium (Ca) in alkaline soils (Jackman and Black 1951, von Wandruszka 2006, Gerke 2015). Despite being heavily bound in soils, utilization of inositol phosphate by plants via the presence of soil microorganisms has been reported (Richardson et al. 2001). Bacteria and fungi have been shown to produce enzymes that can hydrolyze phytate to release orthophosphate (Richardson and Hadobas 1997, Idriss et al. 2002, Lim et al. 2007, Huang et al. 2009). One class of these enzymes found in bacteria are β-propeller phytases (BPPs), which have been shown to specifically hydrolyze phytate under neutral and alkaline pH conditions (Shin et al. 2001, Mullaney 2007, Shim and Oh 2012). This class has a narrow substrate-specificity towards phytate, and various crystal structures analyses have shown that the BPP phytase active site accommodates Ca-bound phytate (Ca-Phytate) (Kerovuo et al. 2000, Shin et al. 2001, Oh et al. 2004)and some studies suggest that solubilization of the Ca-phytate is not necessary prior to mineralization (Oh et al. 2006, Kim et al. 2010). To-date, the BPP is the only microbial phytase shown to work under alkaline conditions; in addition, the phosphatases PhoD and PhoX have been shown to be active under neutral-alkaline conditions. Recently, PhoD and PhoX have been predicted to contribute to increased phytate mineralization in soil (Ragot et al. 2016, Neal et al. 2017) likely via accommodating lower order *myo*-inositol phosphates (IP_5_ – IP_1_), though their activity towards phytate or Ca-phytate has, to our knowledge, not been explicitly tested. The existence of these enzymes suggests a potential for development of biofertilizers using bacteria adapted to alkaline soil conditions with selective advantages for degrading Ca-phytate and lower order *myo*-inositol phosphates.

In the present study, a soil-based microcosm was developed to study microbial communities participating in soil organic P cycling and provide an isolation platform for future biofertilizer candidates. In order to enrich for competitive soil community microorganisms capability of overcoming the preexisting soil ecology, we perform the enrichment *in solum*, with precipitated Ca-phytate used as a bait substrate. A top soil which had not been exposed to conventional P fertilization for at least a decade was used to increase of the potential for encountering bacterial taxa capable of utilizing complex P forms. These microcosms were used to test the hypotheses that; 1) Ca-phytate hotspots recruit distinct populations compared to regions without Ca-phytate and 2) Ca-phytate hotspots recruit bacterial populations with increased phytase activity, and an increased potential for organic P cycling in general. Amplicon sequencing of the 16S rRNA gene and catabolism fingerprinting using BioLog^®^ confirmed significantly divergent general catabolism profiles and taxonomic differences within the recruited communities, and phytase assay and post-incubation *myo*-inositol phosphate quantification supported increased phytase activity. Quantification of bacterial genes associated with organic P cycling in alkaline soils (*phoX, phoD*, and *BPP*) indicated that phosphatases PhoX and PhoD may play a larger role in phytate mineralization in soil than previously recognized. Collectively, we propose that the microcosm approach represents a promising new method for isolation of bacteria candidates with specific functional traits and brings a critical advantage in addressing the preexisting soil ecology during the selection process.

## 2. Materials and Methods

### 2.1 Soil characterization

The soil was collected from a long-term nutrient depletion trial (LTNDT, N_1_K_1_) deprived of P and potassium (K) fertilizers since 1964 (van der Bom et al. 2017). Soil pH and analysis of orthophosphate content were determined by independent lab analysis in triplicate (OK Laboratorium for Jordbrug, Denmark). Soils were extracted with sodium hydroxide–ethylenediaminetetra-acetic acid (NaOH–EDTA) using standard procedures for nuclear magnetic resonance (NMR) analysis based on those of Doolette et al. (2017). For quantification of P into specific classes the ^31^P NMR spectra were integrated across four broad chemical regions: phosphonate (δ18-22 ppm), orthophosphate and phosphomonester P (δ6.5-3.5 ppm), phosphodiester P (δ2 to −1 ppm) and pyrophosphate (δ-4.5 to −5.5 ppm). Due to several overlapping peaks within the orthophosphate and phosphomonoester region of the ^31^P NMR spectra, the relative concentrations of P species were determined using the spectral integration and deconvolution fitting technique described in McLaren et al. (2015).

### 2.2 Set-up and employment of the Ca-phytate soil microcosm baiting system

A microcosm employing a ‘bait’ source of interest, in this case Ca-phytate, was designed for an *in-situ* soil incubation in order to recruit and identify microorganisms inherently successful under competitive soil conditions and attracted to precipitated phytate. In some treatments, an artificial root exudate nutrient solution was used. Figure 1 visually represents the designed hotspot.

**Figure 1.**
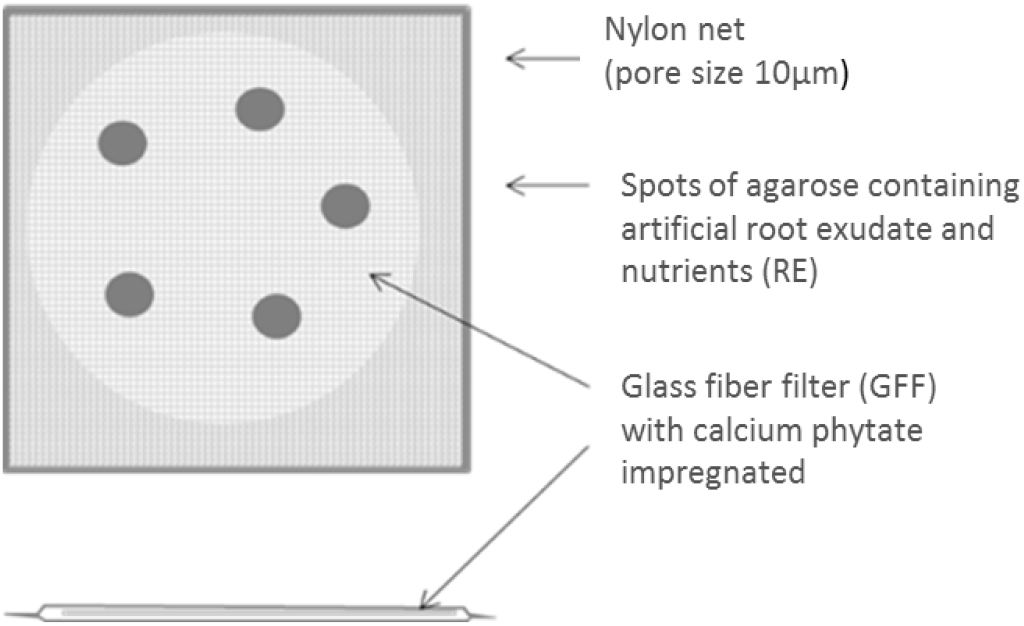
In-soil bait system used for selecting soil bacterial strains attracted to precipitated phytate. The baiting system is composed of a glass fiber filter loaded with experimental and control treatments. Finally, the microcosm is wrapped in sterile mesh and is heat-sealed along all edges in order to exclude soil particles. This example shows the set up for the Ca-IP_6_+RE sample.

The Ca-phytate (Ca-IP_6_) ‘bait’ substrate used was synthesized with modifications based on Hoff-Jøorgensen (1944) *et al*. by adding autoclaved calcium chloride (CaCl_2_) into filter sterilized sodium (Na)-phytate salt (0.2 μm filter) at a 1:6.5 molar ratio and adjusting the pH to 7 using sterile 2 M NaOH. Three wash steps were included in order to eliminate contaminating un-precipitated salts before speed vacuum dehydration. The purity of the phytic acid-Na salt source (BHD biochemical, No 38045) was confirmed via P^31^NMR analysis as well as colorimetric testing to be phytic acid with no contaminating lower order inositols (>99% purity) and negligible free orthophosphate. The resulting Ca-phytate was confirmed via ICP analysis to contain a 1:1 ratio of Ca:P. A volume of 200 μl of a 5% Ca-phytate slurry in water was used to impregnate a 25 mm Whatman™ glass microfiber filter GF/C™ (GFF) and left to dry under sterile conditions. This provided a total amount of 10 mg Ca-phytate on each filter. Two-hundred μl sterile water was used in control hotspots lacking Ca-phytate. An agar solution at ~50°C containing or excluding an artificial root exudate (RE) and nutrient solution was then added dropwise (10 drops of 5 μl per filter) around the GFF filter surface and left to solidify under a laminar flow hood. The RE-nutrient solution consisted of D-glucose (50 mM), D-fructose (50 mM), sucrose (50 mM), succinic acid (25 mM), L-malic acid (25 mM), L-arginine (12.5 mM), L-serine (12.5 mM), L-cysteine (12.5 mM) and nutrients provided as CaCl_2_ (27 mM), NH_4_NO_3_ (30 mM), KCl (67 mM), MgSO_4_ (2 mM), FeSO_4_ (36 μM), MnSO_4_ (45 μM). The treated GFFs were finally wrapped and sealed in 50 μm sterile mesh (Sintab Product AB, fabric 6111-005043) and incubated in soils with water added to 50 % water-holding-capacity (~14 %).

The experimental setup included the following hotspot compositions; i) **Ca-IP_6_**: Ca-phytate treated filters + water agar drops, ii) **RE:** water treated filters + root-exudate agar drops iii) **Ca-IP_6_+RE**: Ca-phytate treated filters + root-exudate agar drops, with the following replicates per treatment; ten GFFs for DNA extraction, six GFFs for BioLog^®^ Eco plates, and three GFFs plus two non-incubated controls for post-incubation Ca-phytate HPLC analysis. The hotspots were completely embedded in LTNDT soil and incubated for 5 days at 20°C. After incubation, hotspots were removed and cut free from the surrounding mesh-bag using flame-sterilized scissors and forceps.

### 2.3 Quantification of *myo*-inositol phosphates recovered from hotspot GFFs by HPLC

Hotspot GFFs retrieved from the microcosm treatments inoculated in soil as well as non-inoculated controls were extracted by shaking for 30 min in 5 mL 0.6M HCl at room temperature. Extracts were filtered using a 0.22 μm filter. Subsequently, the solutions were diluted 1:25 with water and passed through a column containing AG1-X8, 100-200 mesh resin. The column was washed with 10 volumes (the column volume) of water and 10 volumes of 25 mM HCl. Phytate was finally eluted with 5 volumes of 2 M HCl. The eluate was concentrated in a vacuum evaporator to dryness and the residue was dissolved in 1 mL water. Finally, 20 μl of the samples were subject to HPLC (Ultrasep ES 100 RP18; 2 × 250 mm) to determine phytate concentration and presence of lower order *myo*-inositol phosphate derivatives. The column was run at 45°C and 0.2 mL min^−1^ of an eluent consisting of formic acid:methanol:water:tetrabutylammonium hydroxide (44:56:5:1.5 v/v), pH 4.25, as described by Sandberg and Ahderinne (1986). Phytic acid, dodecasodium salt hydrate (Aldrich, 27,432-1) was used as a standard.

### 2.4 Catabolism fingerprinting of the hotspot extracts by BioLog^®^ Eco plates

Samples for BioLog^®^ ECO plate catabolism analysis were performed in triplicate on two hotspot GFFs combined across treatments to obtain sufficient biomass for the assay. Filters were extracted from soil as described above, and the hotspot GFFs were immersed into 11 ml of 0.9% NaCl and shaken at 150 RPM (KS250basic, IKA labortechnik) for 1 hour at room temperature with intermittent vortexing. A subsample was taken and nystatin was added to a final concentration of 50 μg ml^−1^, before aliquoting onto BioLog^®^ Eco plates in triplicate (one ECO plate per sample, 100μl per well). The BioLog^®^ECO plates were incubated at 15°C (to mimic Danish spring time temperatures) in the dark and tetrazolium violet oxidation (color development) was measured at 590nm using an Epoch (BioTek) plate reader every 6-12 hours for 240 hours (10 days). Several time points were analyzed in order to follow the color development, however, the time point chosen for presentation and statistical analysis was the reading corresponding to half of the average well color development (AWCD), which was 144 hours for all treatments. Absorbance levels were normalized to the control well containing only microcosm extract and water. Data was visualized by principal component analysis (PCA). Statistical relevance was tested using a one-way ANOSIM (non-parametric) employing Euclidean distance measures. Post-hoc ANOSIM pairwise comparisons identified specific treatments contributing to significance at *P* < 0.05.

### 2.5 Phytase activity of the hotspot extracts -

The remaining sample of the above 0.9% NaCl microcosm extraction was used for measuring phytase activity; hence, the remaining volume after subsampling for BioLog analysis and the GFF itself were subjected to mechanical lysis to recover the entire microbial biomass associated with the microcosm. This was considered as a cell-associated enzyme fraction. The phytase assay used here differed from standard phytate assays, as it utilized precipitated Ca-phytate at pH 7, whereas the standard phytase assay utilize soluble Na-phytate at pH 5 (Jukka K. Heinonen 1981). The Ca-phytate was prepared by adding filter-sterilized *myo*-inositol hexa*kis*phosphate dodecasodium salt (Na-IHP; Sigma P-0810) suspended in distilled water to sterile 1.5M CaCl_2_, and bringing the pH up to 7 using sterile NaOH. The resulting precipitate was washed in sterile distilled water and suspended to an appropriate concentration for immediate use. The hotspot GFFs were suspended in a total of 1 mL of 0.9% NaCl in Lysing Matrix E tubes containing glass beads (MP Biomedicals) and beaten 3 times at 1 oscillation/s frequency (FastPrep-24, MP Biomedicals). The samples were then centrifuged at 10,000 x *g* for 10 min at 4 °C and the aqueous fraction used immediately in phytase assays. Orthophosphate released in the phytase assays was measured in a 96-well microtiter plate using malachite green (MG) colorimetry as described by D’Angelo et al. (2001). Briefly, 30 μl of MG reagent (1 g L^−1^ polyvinylalcohol, 1.2 g L^−1^ H_3_BO_3_, 34.66 g L^−1^ (NH_4_)_2_MoO_4_, 476 ml L^−1^ 10N H_2_SO_4_, 0.229 g L^−1^ malachite green) was added to each well containing 240 μL of the crude enzyme extract. The assay was incubated for 24 hours at room temperature as described by Tang et al. (2006). Assay conditions included the presence and absence of Ca-phytate salt precipitate (20 mM Ca-IP_6_ (as described above) and MOPS (20 mM) buffered at pH 7.2. Absorbance was measured at 620 nm after allowing 30 minutes color development. Orthophosphate standards from 0 to 40 μM were used as a standard curve. Technical duplicates were run on all samples and controls. Extracts assayed in the absence of substrate were included to account for residual orthophosphate or phytate degradation under the assay conditions and were subtracted from the experimental values. The resulting data were normalized to bacterial 16S rRNA gene copy number as determined by qPCR (as outlined below).

### 2.6 Quantitative PCR (qPCR) of bacterial 16S rRNA and functional genes

Two GFFs were collected from each treatment and washed in a 5 mL aliquot of 0.9 % NaCl as described above. Quintuplicate samples were then taken from the pooled extract for DNA extraction. A phenol-chloroform protocol for DNA extraction was employed as described previously (Nicolaisen et al. 2008), with the exception that all samples were incubated overnight at 4 °C during PEG precipitation. Primer sequences used for quantification of the bacterial 16S rRNA gene, the alkaline phosphatase genes *phoD* and *phoX;* and the β-propeller phytase gene (*BPP*) are listed in supplementary materials (Table S1). Absolute quantification was performed using plasmid-based standard curves developed by Hao et al. (2020). All reactions were performed on the AriaMX^®^ using the Brilliant III UltraFast QPCR Master Mix SYBR^®^ Green (Agilent technologies, USA). Each 25 μL reaction contained 2 μL of the undiluted microcosm DNA extract, 1 μg μL^−1^ BSA (New England Biolabs ^®^ Inc., UK), 1× mastermix and 0.4 μM primer. Thermal profile for *BPP* primers was carried out as follows: 3 minutes at 95°C followed by 5 cycles touch-down starting at 65°C and decreasing 0.4°c per cycle, after which 35 cycles of 30 s at 95 °C, 30 s at 63 °C, 30 s at 72 °C, and 10 s at 82 °C were carried out. The thermal profile for all other primers was as follows: 3 minutes at 95 °C followed by 40 cycles of 20s at 95 °C, 30 s at 58 °C. Fluorescence was measured at the end of the final step in each recurring cycle. A melting curve analysis was performed in each run to assist in quality control checking for each reaction. The absolute abundance of specific functional genes was calculated based on standard curves generated by 10-fold serial dilutions of standard plasmids and was normalized to the 16S rRNA gene (*i.e*. gene copy number/ 16S rRNA gene copy number; Hao et al. (2020). The efficiency of the standard curves ranged from 76 % to 83 %, indicating a potential underestimation in this assay. All samples were run using 5 independent biological replicates extracted in triplicate. For absolute numbers (*phoX, PhoD, BPP*) a Kruskal-Wallis test for non-normally distributed data was used, followed by Mann-Whitney pairwise comparisons. For normalized data (*i.e*. gene/16S rRNA copy number), a square root transformation (sqrt(value+1)) or BoxCox transformation (Y^−λ^=1/Y^λ^) was performed to achieve normality prior to performing one-way ANOVA. The *P*-values reported are either as Fishers ANOVA, where equal variance is followed, or by a Welche’s ANOVA in the case of unequal variance as tested by Levene’s test for homogeneity of variance based on means. Either a Tukey’s or Games-Howell (in the case of unequal variance) pairwise comparison was used *post-hoc* to identify specific treatments contributing to the ANOVA significance.

### 2.7 Taxonomic identification and comparative analysis by 16S rRNA amplicon sequencing

All samples were sequenced by Macrogen Inc. (Seoul, Rep. of Korea) using aliquots of the above-mentioned DNA extracts. In short, the variable region V3-V4 of the 16S rRNA gene was amplified by PCR using the primer pair 341F (5’-CCTAYGGGGRBGCASCAG-3’) and 806R (5’-GGACTACNNGGGTATCTAAT-3’) and sequenced using a Paired-end Illumina MiSeq platform. A total of 6,087,974 16S rRNA gene sequence reads were obtained. After quality filtering and removal of adaptors and ambiguous nucleotides, 2,686,025 merged pairs were obtained with an average PHRED score of 45. Sequences were analyzed using CLC Genomics Workbench with the Microbial Genomics Module (Qiagen). Sequence adaptors were removed and trimming was performed to exclude low quality DNA (quality score limit 0.05) and sequences with ambiguities above 2. Reads less than 5 nucleotides were discarded before merging forward and reverse amplicon reads with default values and standardizing sequence lengths. The partial 16S rRNA gene sequences were then clustered and assigned to operational taxonomic units (OTUs) with a 97% similarity using the Greengenes database (DeSantis et al., 2006) with allowance of new OTU creation. During OTU clustering suspected chimeras were removed and finally, taxonomy including chloroplast or mitochondria were filtered from the sequence list. The sequence reads clustered into 1,434 known OTUs, as well as 550 *de novo* OTUs; yielding a total of 1,984 OTUs. Rarefaction analysis supports that the sequencing depth captured sufficient diversity of the bacterial communities in the microcosm and bulk soil samples, as seen by apparent asymptote in the rarefaction curves (Supplementary Figure S1). The α-diversity and Principal Coordinates Analysis (PCoA) were performed with the software PAST v.2.17 using Bray–Curtis dissimilarity matrices at an OTU level. Furthermore, pairwise similarity percentages (SIMPER) were used to determine the taxa contribution to the average dissimilarity between treatments (Bray-Curtis calculated) (Clarke 1993) and were ultimately used to create a Venn diagram; using VENNY (Oliveros 2007). The calculated α-diversity, represented by Shannon, and Chao-1 estimations, was compared using ANOVA and *post-hoc* Tukey test. Rarefication to the lowest read number using RStudio (V 1.1.456) software was performed before testing for statistical significance of the overall difference in microbial community composition (β-diversity) by one-way NP-MANOVA (PERMANOVA) on Bray-Curtis dissimilarity matrix with 9999 permutations in PAST v.2.17.

### 2.8 Accession numbers

Sequences were deposited in the European Nucleotide Archive (ENA) database under the study accession number PRJEB39274.

## 3. Results

### 3.1 P content of soil used for the microcosm

P^31^NMR analysis of the LTNDT soil showed a total amount of extractable P of 293 mg/kg. Organic P accounted for approximately half of the total P and consisted primarily of α- and β-glycerophosphate (likely derived from the alkaline hydrolysis of lipids in the soil sample), inositol phosphates as well as uncharacterized monoester P, whereas no phosphonate or diesters were detected. Of the inositol phosphates, *myo*-inositol hexa*kis*phosphate (phytate or IP_6_) accounted for 27 mg/kg, corresponding to 9.3% of the total soil P. The extractable Olsen P and pH (CaCl_2_) for the soil were found to be 6.5 ± 1.5 mg/kg and pH 6.13 ± 0.04, respectively.

### 3.2 Recovery of *myo*-inositol phosphate (phytate) from hotspots

Extraction from the filter hotspots and quantification by HPLC represented a *myo*-inositol phosphate recovery efficiency of 95.5%, compared with the non-incubated GFFs as reference samples. The soil-incubated microcosms containing Ca-phytate plus root exudate (Ca-IP_6_+RE) showed a significant decrease in HPLC-detectable *myo*-inositol phosphate with quantities averaging 1.76 mM (83% of initial, *post-hoc* Mann-Whitney *P* < 2.4 × 10^−4^, **Table 1**), whereas the soil-incubated Ca-phytate hotspot (Ca-IP_6_) alone showed no detectable loss of *myo*-inositol phosphate.

**Table 1.** *Myo*-Inositol phytate substrate remaining on the microcosm GFF pre and post in-soil incubation. Data obtained from soil-incubated samples were based on biological triplicates and non-incubated samples were tested in biological duplicates. Statistical relevance was tested using the non-parametrical Kruskal-Wallis test. Microcosms treatments were as follows: calcium phytate alone (Ca-IP_6_), root-exudate alone (RE), and calcium phytate combined with root-exudate (Ca-IP_6_+RE).

| Microcosm treatment | Soil incubation | $IP_6$ [mM] | SEM (n = 3) |
| --- | --- | --- | --- |
| Ca-$IP_6$ | no | 2.115 | 0.014 |
| RE | no | nd | - |
| Ca-IP<sub>6</sub> | yes | 2.126 | 0.025 |
| RE | yes | nd | - |
| Ca-IP<sub>6</sub>+RE | yes | 1.765 * | 0.041 |

\* Significant decrease in IP<sub>6</sub> detected when compared to non-soil incubated and incubated Ca-IP<sub>6</sub> controls (*post-hoc* Mann-Whitney; $P < 2.4 \times 10^{-4}$ ), nd = non-detectable

Soil incubated filters containing only RE yielded no detectable substrate regardless of their incubation status, indicating no soil-originating contamination of Ca-IP_6_ presence. No lower order *myo*-inositol phosphate products were detected in any of the treatments, despite the capacity of the HPLC assay being able to detect lower order *myo*-inositol phosphates IP_5_ – IP_3_.

### 3.3 Orthophosphate liberation by crude enzyme hotspot extracts

Crude activity of the enzyme extracts from the microcosms harboring the Ca-phytate hotspot showed release of orthophosphate that was in the order of 23.6 ± 13.2 nmol and 54.4 ± 36.5 nmol of P from the Ca-IP_6_ and the Ca-IP_6_+RE hotspots respectively. No detection of released orthophosphate was seen in the crude extract acquired from the RE microcosm. To account for the varied bacterial recruitment across the treatments, these values were normalized to 16S rRNA gene copy numbers (i.e., nmol P /16S rRNA gene copy number) showing comparable P release per 16s rRNA (**Figure 2**). Whilst no statistical difference was seen between treatments (Ca-IP_6_ and Ca-IP_6_+RE), it is apparent, that no detectable orthophosphate was released from the crude extracts originating from microcosms lacking Ca-phytate (RE).

**Figure 2.**
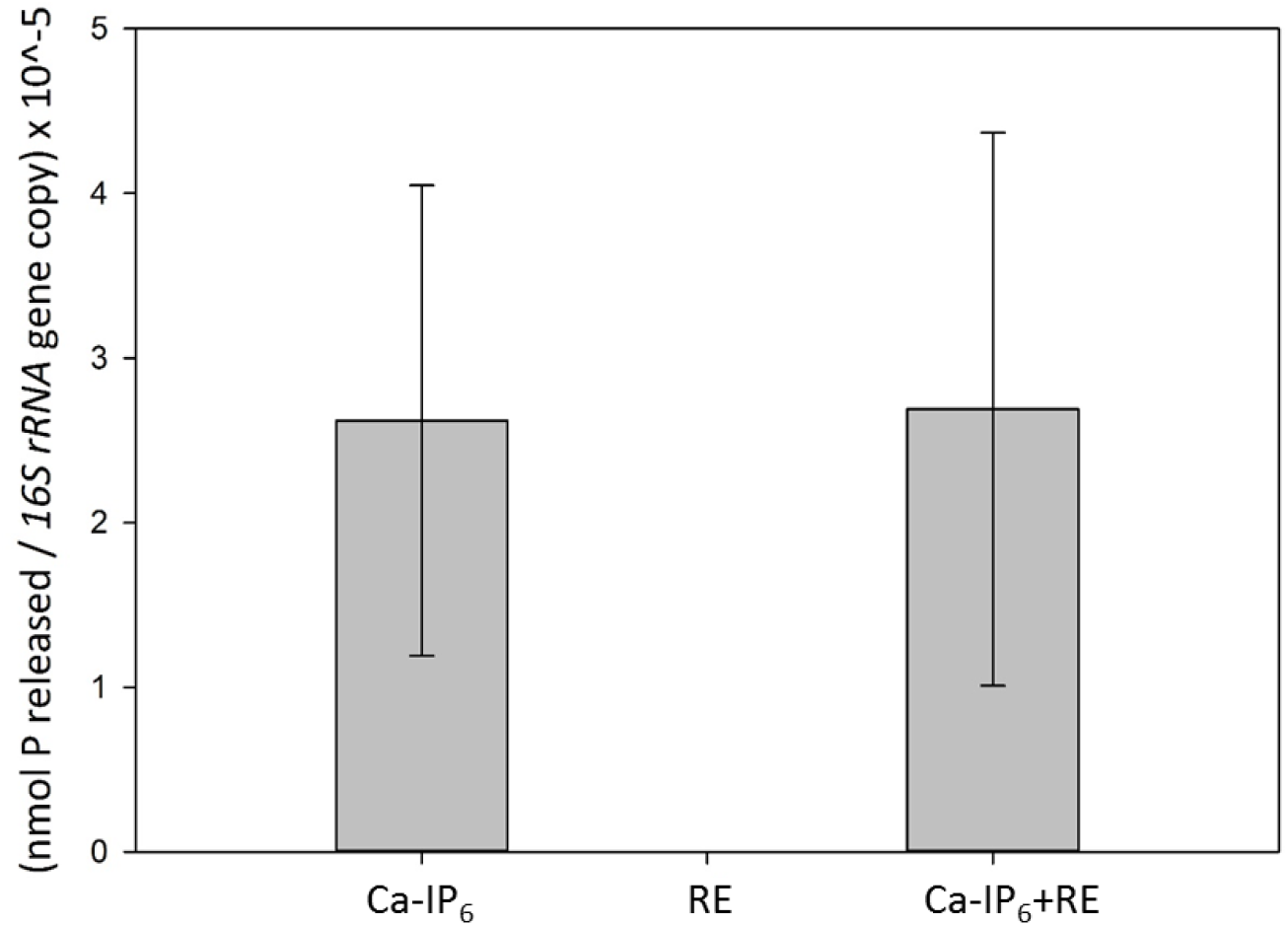
Phytase assay on treatments showing released orthophosphate normalized to 16S rRNA gene copy number. Values are calculated per GFF extraction using crude enzyme extracts of cell-associated materials in triplicate. The 16S rRNA copy numbers were determined as described in materials and methods. No activity was detected in the microcosm crude extract containing only root exudate (RE). Microcosms treatments were: calcium phytate alone (Ca-IP_6_), root-exudate alone (RE), and calcium phytate combined with root-exudate (Ca-IP_6_+RE). Error bars are reported in SEM.

### 3.4 Quantification of hotspot-associated bacteria and quantification of functional genes involved in organic P cycling

In order to understand the population size in the bulk soil and between treatments, the total extractable 16S rRNA genes were quantified by qPCR. The bulk soil yielded 3.53 × 10^7^ ± 0.12 × 10^7^ copies per gram of soil (Supplementary Figure S2). Different total number of bacteria were recruited from the various treatments (Kruskal-Wallis, *P* = 2.6 × 10^−7^, **Figure 3**).

**Figure 3.**
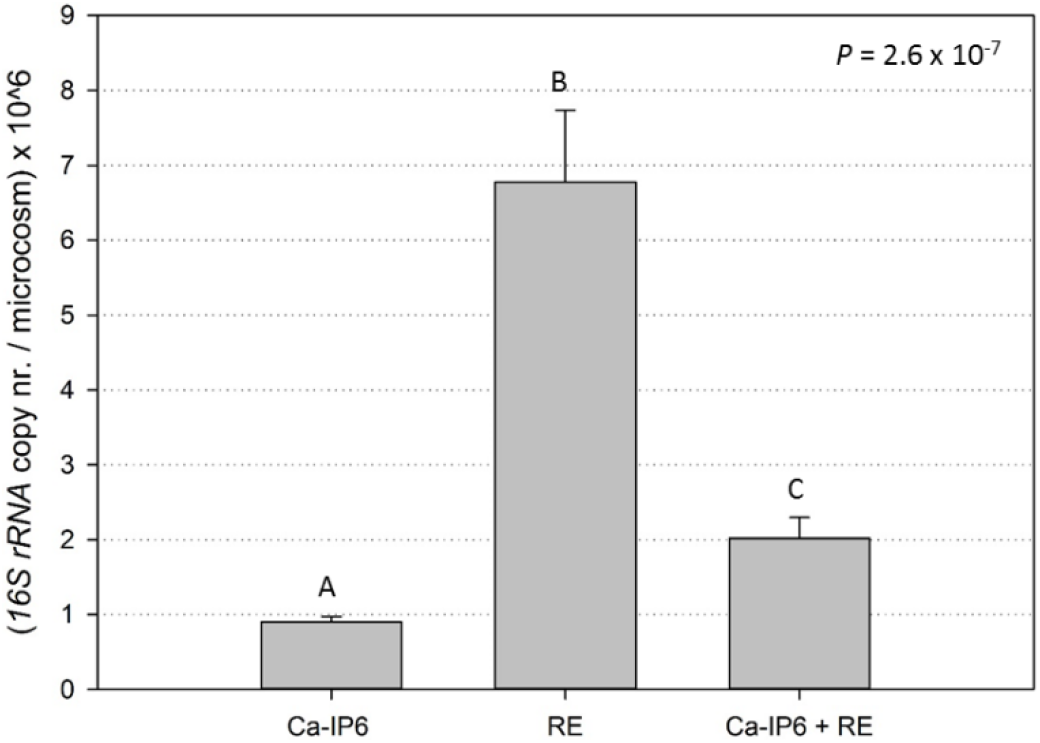
16S rRNA gene copy numbers per microcosm. Microcosms treatments were: calcium phytate alone (Ca-IP_6_), root-exudate alone (RE), calcium phytate combined with root-exudate (Ca-IP_6_ + RE). The P-value shown is generated from the Kruskal-Wallis test and letters denote pairwise Mann-Whitney results. Error bars are shown in SEM.

The RE hotspot recruited more 16S rRNA gene copies than the hotspots containing Ca-phytate (*P* < 6.8 × 10^−5^). Addition of root exudate to the Ca-phytate embedded GFF (*i.e*. Ca-IP_6_+RE) resulted in significantly more gene copies than hotspots containing only Ca-phytate (*i.e*. Ca-IP_6_, *P* = 0.002) alone, however, also significantly less than root exudate on its own (*P* = 6.8 × 10^−5^).

Based on absolute gene copy numbers, *phoD* was most abundant followed by *phoX*, and then BPP (Figure 4). The total number of phosphatase gene copies (*i.e*. *phoD* + *phoX* + *BPP*) are highest in the RE treatment (5230 copies ± 1048) as compared to Ca-IP_6_+RE (3676 ± 211) and Ca-IP_6_ (2233 ± 535)(errors reported in SEM, Figure 4.) The *phoD, phoX*, and *BPP* genes were also measured per gram bulk soil (supplementary data, Figure S2), however, are not comparable to the absolute data from the microcosm, which is extracted from GFFs.

**Figure 4.**
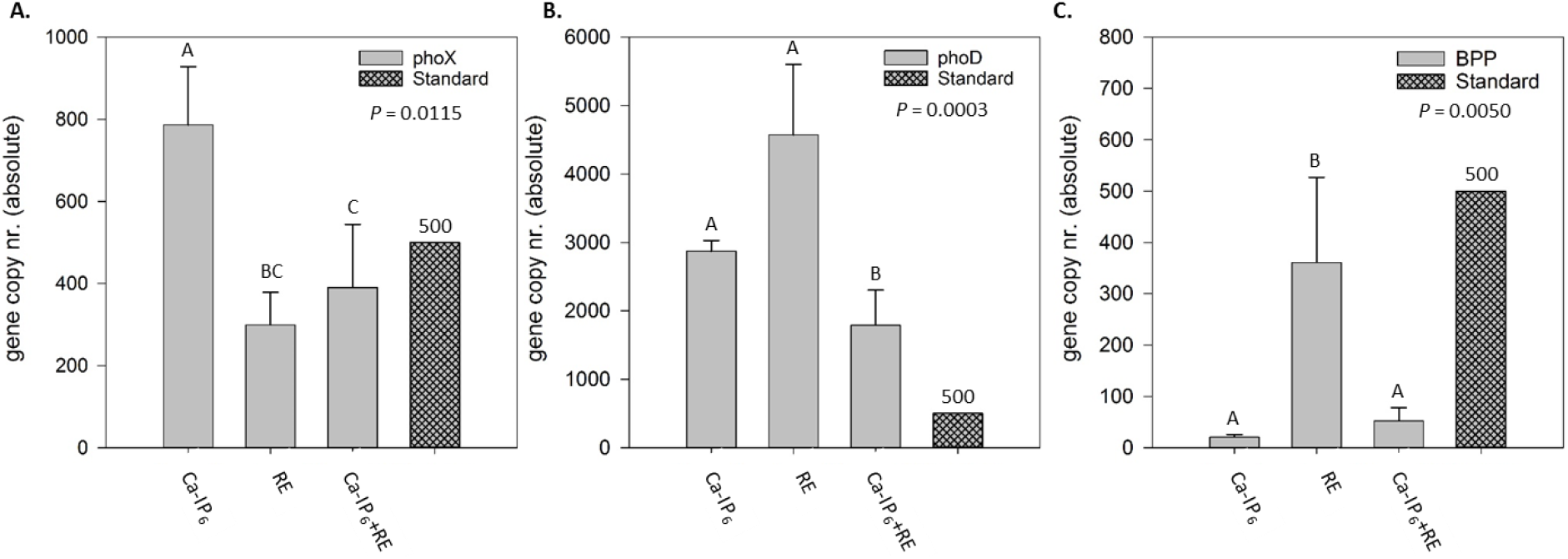
Absolute abundance of genes relating to organic phosphorus seen in treatments. A. *phoX* = alkaline phosphatase, B. *phoD* = alkaline phosphatase, C. BPP = beta propeller phytase. Microcosms treatments were: calcium phytate alone (Ca-IP_6_), root-exudate alone (RE), calcium phytate combined with root-exudate (Ca-IP_6_+RE).. Error is shown in SEM and statistical comparisons are done using Kruskal-Wallis with Mann-Whitney post-hoc pairwise evaluations. A shaded standard bar is included to assist in drawing comparisons between graphs.

The relative abundance of *phoX, phoD*, and BPP were similarly calculated as gene copy per 16S rRNA gene copy number across all treatments as well as in bulk soil (Figure 5). Compared to bulk soil, specific enrichment of *PhoX* and *PhoD* was evident in the Ca-phytate hotspot, whereas gene abundance was decreased in the treatments with RE (Games-Howell and Tukey post-hoc respectively, *P* < 0.04). No difference in relative abundance of BPP occurred across any hotspot treatment compared to bulk soil (*P* = 0.24, Welch’s F test).

**Figure 5.**
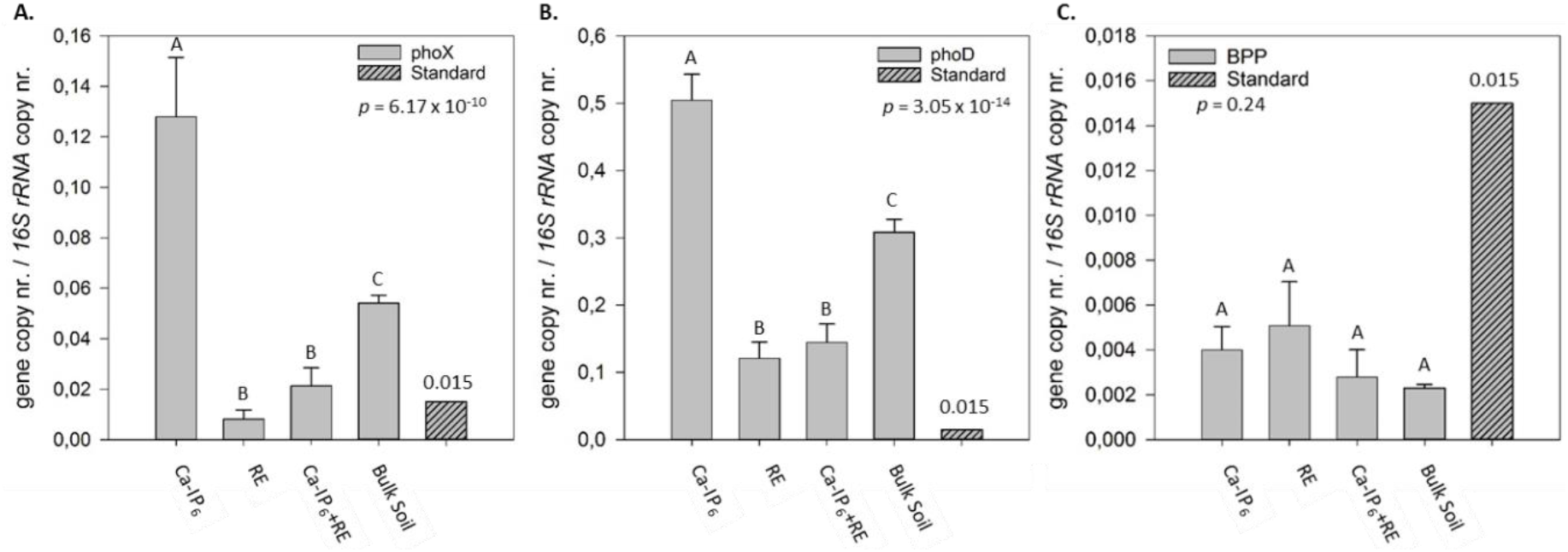
Relative abundance of genes relating to organic phosphorus seen in treatments and bulk soil normalized to 16S rRNA copy number. *phoX* = alkaline phosphatase, *phoD* = alkaline phosphatase, BPP = beta propeller phytase. Microcosms treatments were: calcium phytate alone (Ca-IP_6_), root-exudate alone (RE), calcium phytate combined with root-exudate (Ca-IP_6_+RE), and bulk soil. Error is shown in SEM and statistical comparisons are done using Fisher and Welches ANOVA with appropriate post-hoc pairwise evaluations. A shaded standard bar is included to assist in drawing comparisons between graphs.

### 3.5 Catabolism fingerprinting of bacterial populations using BioLog^®^Eco plate

The hotspot treatment had a significant effect on community composition as assessed by the BioLog^®^Eco plates at all time points tested (P < 0.0001, ANOSIM, Euclidean distance). However, the time point corresponding to half of the average well color development (AWCD) was 6 days (144 hours) and was used for statistical analysis and data comparison. Pairwise ANOSIM comparisons indicated that the response between treatments differed with the following associated *P* values; Ca-IP_6_ and RE hotspots; *P* < 0.001, RE and Ca-IP_6_+RE; *P* = 0.015, and Ca-IP_6_ and Ca-IP_6_+RE; *P* = 0.051 (Figure 6). The metabolic fingerprint indicated that the Ca-IP_6_+RE treatment was between Ca-IP_6_ and RE.

**Figure 6.**
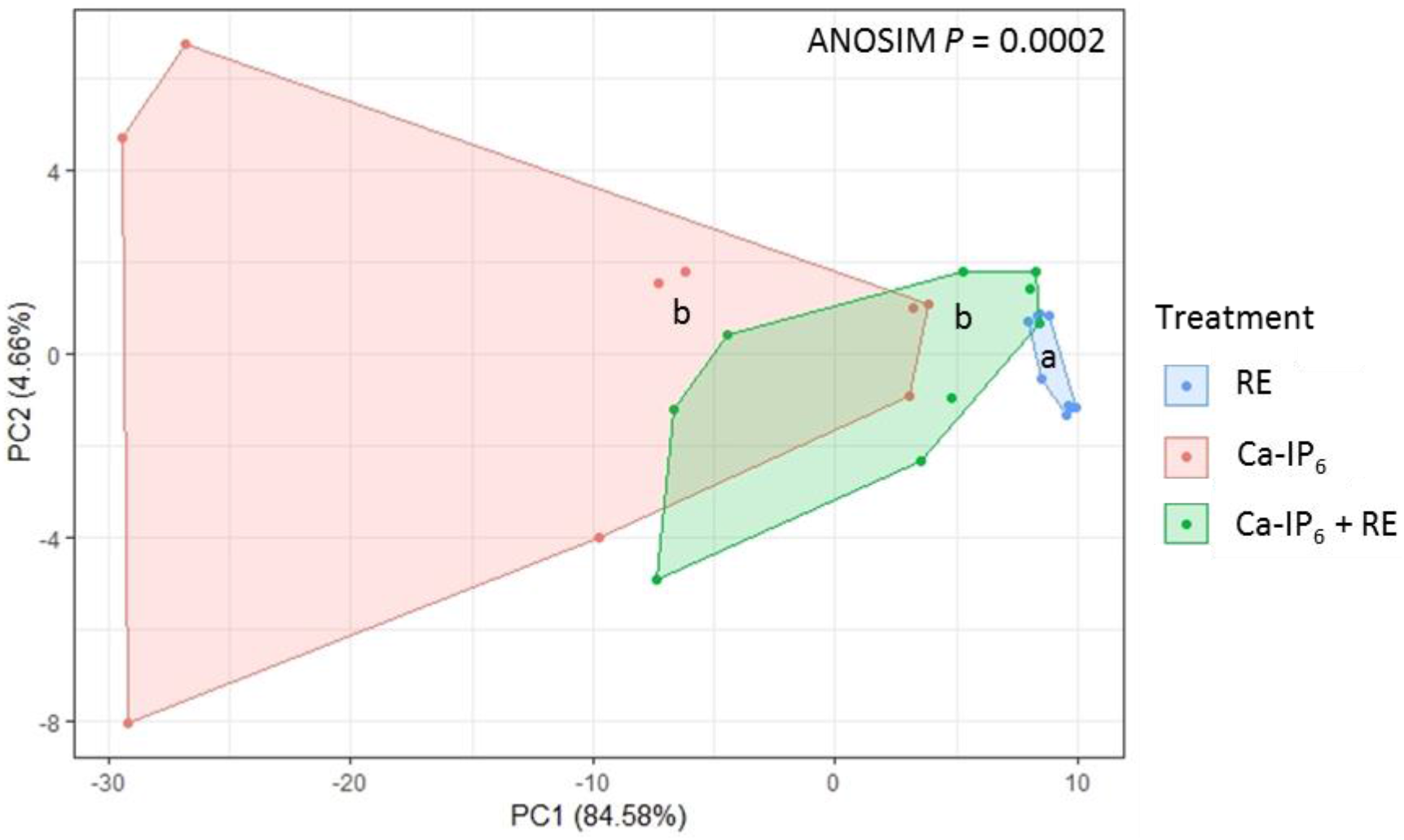
BioLog ECO plate data represented in a principle components analysis (PCA). The shaded areas represent convex hulls enclosing assigned treatment points in the Euclidian plane. Microcosms treatments were: calcium phytate alone (Ca-IP_6_), root-exudate alone (RE), and calcium phytate combined with root-exudate (Ca-IP_6_+RE). A blank microcosm control was not included as insufficient biomass was obtained for the assay.

### 3.6 Diversity comparison between microcosm treatments

Evaluation of α-diversity using a one-way ANOVA confirmed statistically significant effects of hotspot conditions by both Shannon and Chao-1 diversity measures (*P* < 0.0001). *Post-hoc* analysis showed that all microcosm microbial communities had significantly lower α-diversity as compared to the bulk soil microbial communities (P = 0.0002, Tukey, Figure 7A and B). In addition, the RE treatment showed significantly lower α-diversity than the Ca-IP_6_ treatment according to the Chao-1 index (P = 0.029, Tukey, Figure 7A) and Shannon index (P = 0.026, Tukey, Figure 7B), whereas the Ca-IP_6_+RE hotspots revealed an intermediate level of α-diversity.

**Figure 7.**
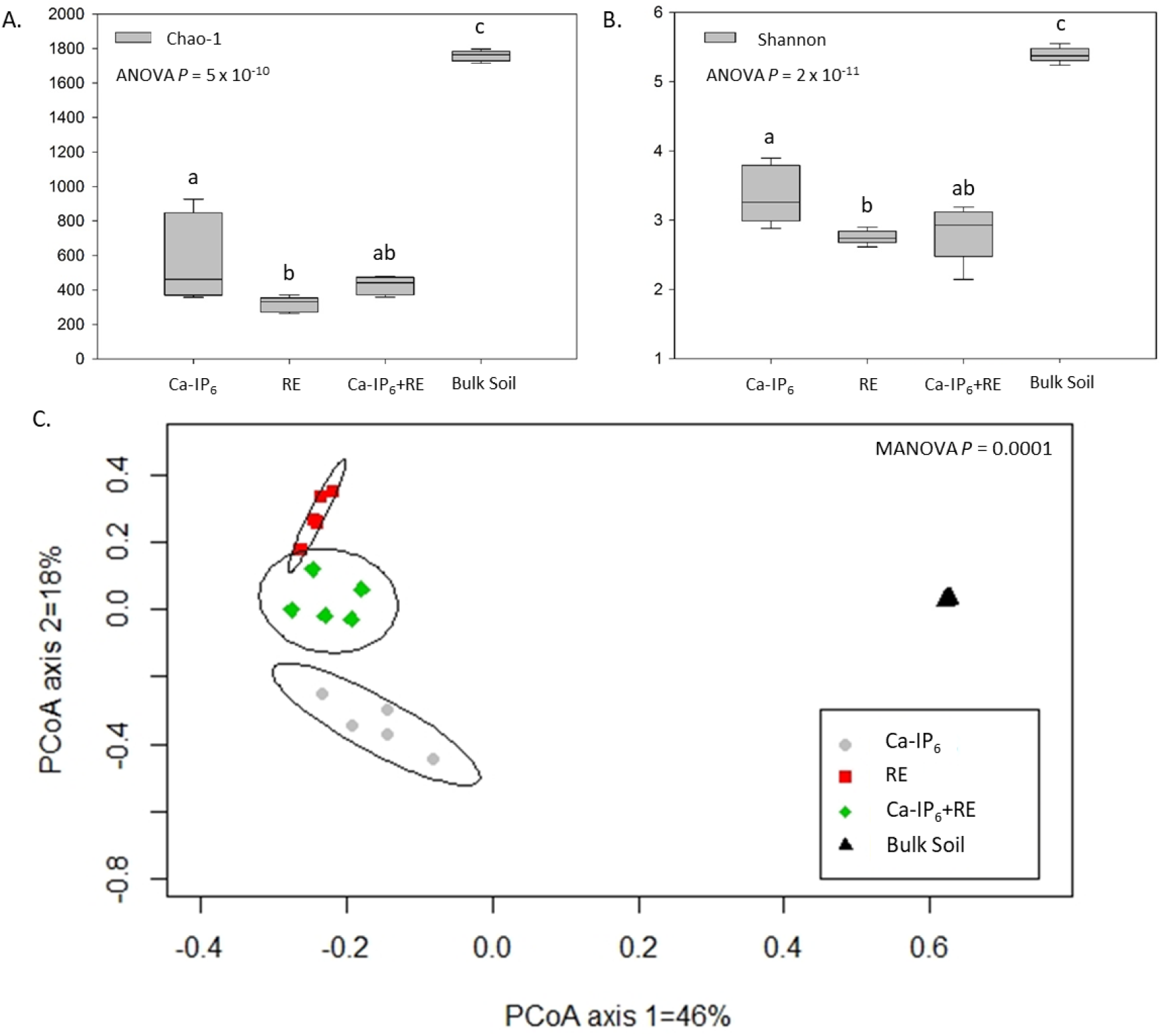
Diversity analysis of microcosm microbial communities. A) Shannon α-diversity and B) Chao-1 α-diversity between microcosm treatments with differences (indicated by different letters) determined using a two-tailed T-test. C. β-diversity across treatment hotspots shown by Principal coordinate analysis (PCoA) of OTUs originating from the microcosm treatments and bulk soil. Treatments were: calcium phytate alone (Ca-IP_6_), root-exudate alone (RE), calcium phytate combined with root-exudate (Ca-IP_6_+RE), and bulk soil. A Bray-Curtis dissimilarity matrix was applied for ordination of the normalized (relative abundance) and rarefied data. Ellipses drawn represent a standard deviation of the data based on 95% confidence interval.

Evaluation of the β-diversity showed significant differences between hotspot treatments as well as bulk soil, as visualized by PCA (Figure 7C). The non-parametrical MANOVA analysis based on Bray-Curtis dissimilarity yielded a *P*-value of 0.0001, with all pairwise comparisons being significantly different (*P* < 0.05) when tested at an OTU level. In the bulk soil, the most abundant phyla were Proteobacteria (43%), Actinobacteria (16%), Firmicutes (15%) and Verrucomicrobia (12%). Acidobacteria, Gemmatimonadetes, Chloroflexi, Planctomycetes, and Bacteroidetes each represented 1 to 4% of the obtained sequences, while the remaining phyla detected each represented less than 0.5% (Supplementary Table 2). Hotspot communities recruited from 53% to 71% OTUs matching the Actinobacteria phylum, followed by the Proteobacteria phylum (22 to 25%). The phylum Firmicutes was represented highest in Ca-IP_6_+RE (24%), followed by RE (12%), and Ca-phytate (4%). Aside from Bacteroidetes being highly represented in RE (7%), all other phyla were represented at levels less than 0.5% (Figure 8A).

**Figure 8.**
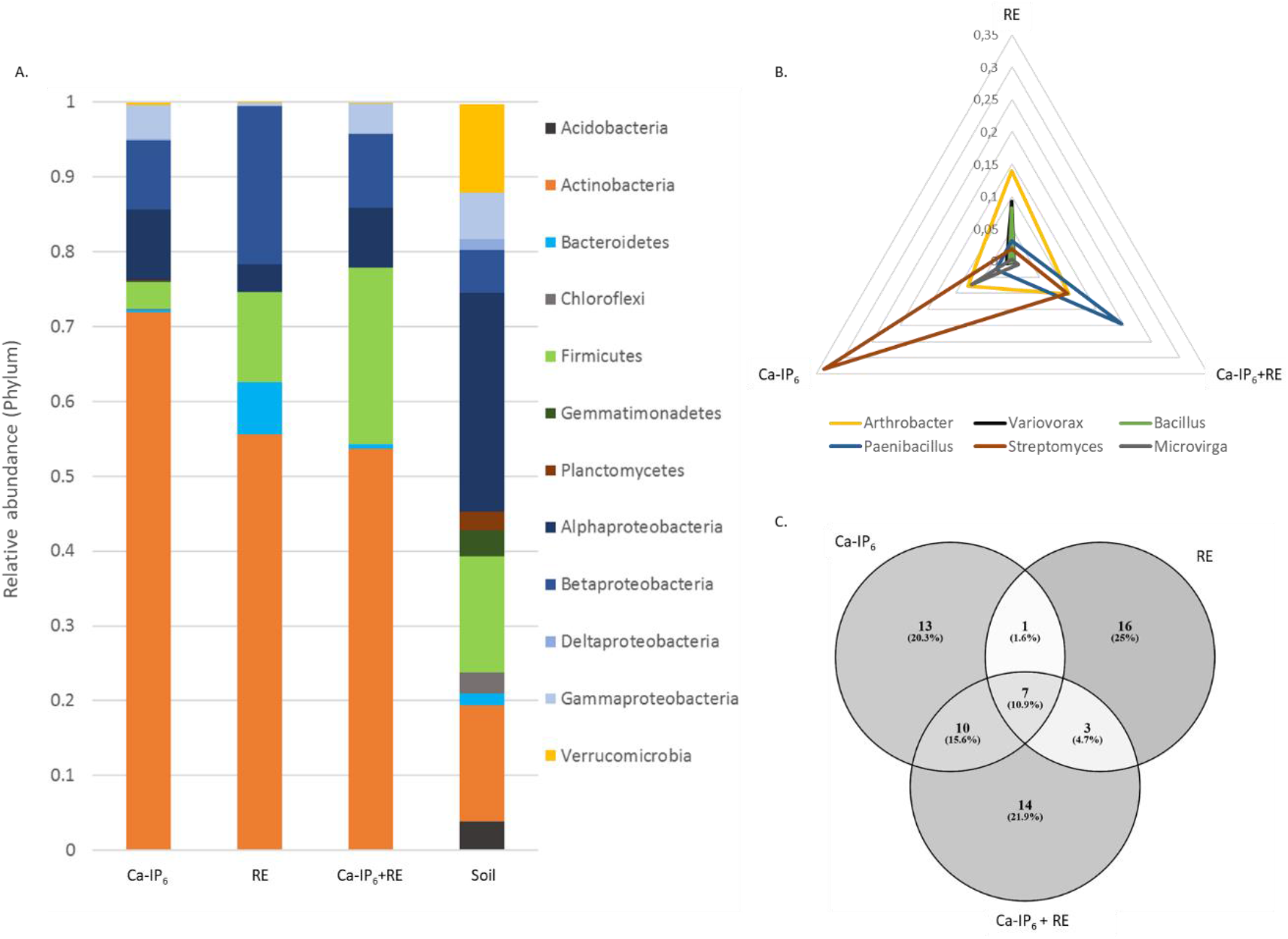
Community composition based on 16S sequence analysis and OTU similarity across microcosm hotspot treatments. A) Stacked bar chart displaying the treatment profiles at a phyla level. OTUs were identified at a 97% similarity level with families representing more than 0.5% abundance being shown. B) Radar plot displaying the top three most abundant assigned genera based on OTUs in each microcosm treatment. C) Venn diagram showing relevant shared and individual OTUs (and total percentage) per hotspot treatment. Only OTUs representing greater than 0.5% abundance were chosen for comparisons. Microcosms treatments were: calcium phytate alone (Ca-IP_6_), root-exudate alone (RE), calcium phytate combined with root-exudate (Ca-IP_6_+RE), and bulk soil.

When looking specifically at the most abundant genera selected within each treatment, preferential selection of *Bacillus* and *Variovorax* was evident in the RE treatment, *Streptomyces* and *Variovorax* in the Ca-IP_6_ treatment, and *Paenibacillus* and *Streptomyces* in the Ca-IP_6_+RE treatment, and relatively equal selection of *Arthrobacter* amongst all treatments (Figure 8B). All OTUs amounting to more than 0.5% total sample contribution were used in a Venn diagram to reveal shared and unique OTUs recruited by the distinct hotspot conditions (Figure 8C). The Ca-IP_6_ and Ca-IP_6_+RE microcosms recruited 13 and 14 unique OTUs, respectively (*i.e*. OTUs that are not recruited in other treatments at an abundance over 0.5%), while 10 OTUs were shared between the treatments (*i.e*. OTUs that were detected in both treatments). Of these 10 shared OTUs that were recruited by the Ca-IP_6_+RE and the Ca-IP_6_ hotspots, eight were classified within the Actinobacteria, the majority within the genus *Streptomyces* (Table 2). To exemplify the quantitative differences between sample treatments, the fold increase of Ca-IP_6_ OTUs relative to the RE-control and the soil community is indicted (Table 2).

**Table 2.**
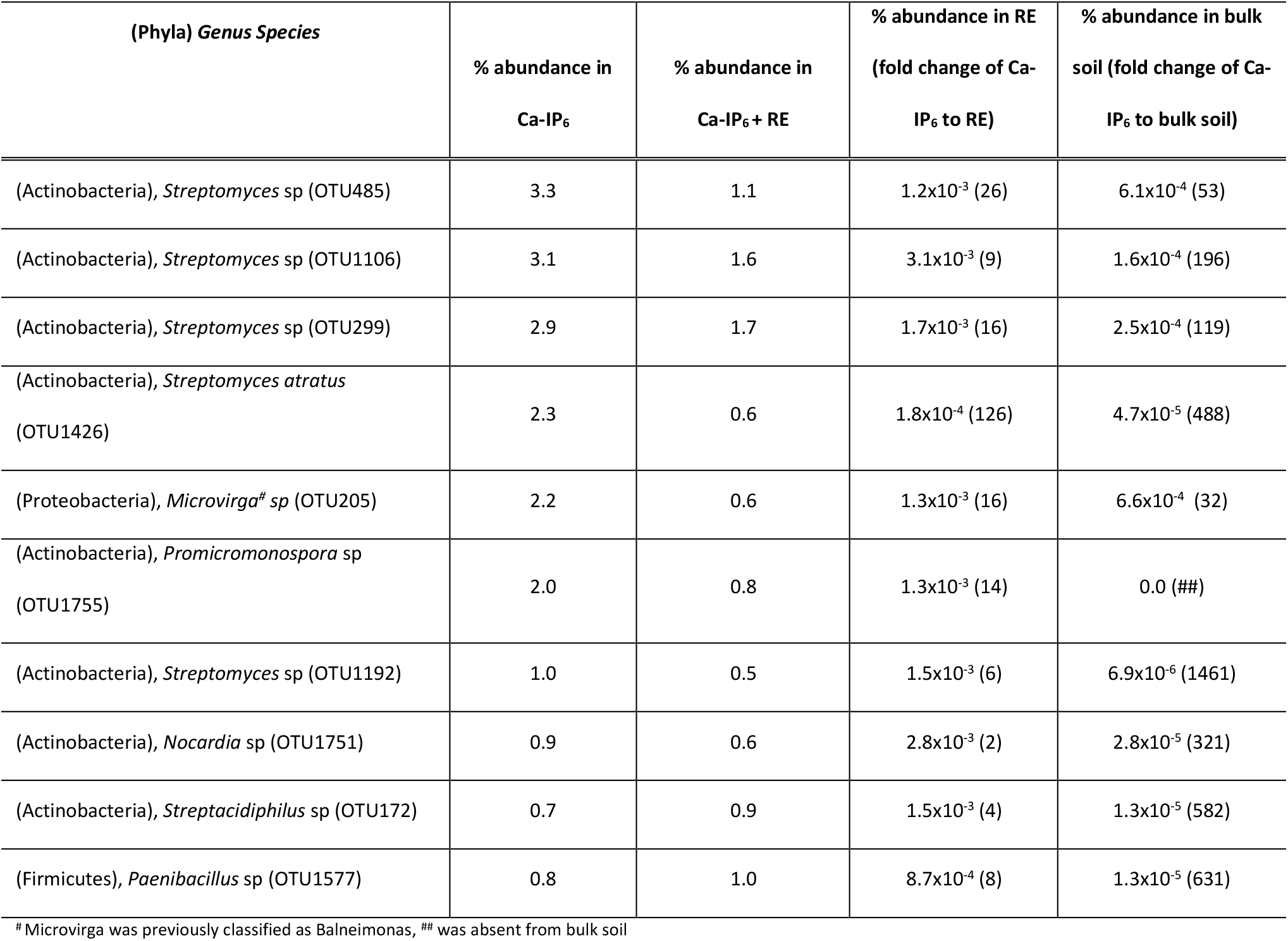
High abundance OTUs shared between Ca-phytate and Ca-phytate + RE treatments. Data is derived from the Venn diagram (Figure 6C) with OTUs having an abundance of higher than 0.5% in Ca-IP_6_ and Ca-IP_6_+RE samples and being absent or lower than 0.5% abundance in RE and bulk soil samples (note that OTUs that are completely absent in RE or bulk soil cannot yield a fold change value)

## 4. Discussion

In the present study, the baiting approach was developed and used to reveal more general knowledge on the ecology of soil microbial communities with potential for organic P mineralization in soil. With the long-term perspective of isolating biofertilizer strains for use in agriculture with improved ability to function in natural soil settings, parameters for the microcosm were thoughtfully selected. The soil used for the microcosm setup contained significant amounts of phytate but low available P, previously shown to be preferred conditions for phytase action (Ramírez and Kloepper 2010). An artificial root exudate was selected to simulate rhizosphere conditions and provide an extra carbon (energy) source beside the Ca-phytate. A 50 μm mesh net was wrapped around the microcosm to create distance between the bait substrate and soil to select for mobile bacteria and keep the filter free from debris. Finally, Ca-phytate was chosen as a relevant “bait” substrate since it is a predicted form of organic P in neutral to alkaline agricultural soils (Jackman and Black 1951, von Wandruszka 2006).

To test if the recruited populations showed any activity towards the bait substrate, breakdown of Ca-phytate was indirectly assessed by HPLC quantification of remaining *myo*-inositol phytate post-incubation and released orthophosphate was quantified from each treatments’ crude enzyme extract in a modified phytase assay. An overwhelming majority of published phytase assays are buffered at pH 5 and utilize a soluble Na-phytate, however, the modified phytase assay used here is buffered at pH 7.2 and utilizes a precipitated Ca-phytate (Engelen et al. 1994, Tang et al. 2006). Using precipitated Ca-phytate, we aim to better mimic the bio-availability challenges of persistent insoluble Ca-phytate salts in alkaline soils. Using a precipitated substrate means that the enzymes cannot not act with maximum velocity (Michealis-Menten kinetics), since the concentration of IP_6_ available to the enzymes is quite low in the reaction mixture. However, despite the added substrate availability challenge, orthophosphate release from Ca-phytate is detectable from both hotspots employing Ca-phytate as a bait substrate (i.e., Ca-IP_6_ and Ca-IP_6_+RE treatments). We consider this an encouraging indication that the population recruited by these hotspots is functionally enriched. Furthermore, in this assay we forfeit extracellular enzyme fractions, as only the fractions physically remaining on the GFF can be removed from soil (i.e., cell-associated extracellular enzymes as well as intracellular enzymes). Therefore, while the values obtained from the phytase assay are low in comparison to other published reports (Richardson and Hadobas 1997, Giles et al. 2018), the above-listed limitations of the modified phytase assay would substantiate this. We consider the limitations of the modified assay a reasonable trade-off for a more representative insight to soil biochemistry surrounding organic P cycling.

While the phytase assay shows orthophosphate release in both Ca-IP_6_-containing hotspots, the HPLC assay could only detect clearance of *myo*-inositol phosphate (IP_6_) in the Ca-IP_6_+RE hotspot (**Figure 2**, **Table 1**). This is likely due to the increased bacterial numbers in Ca-IP_6_+RE, as estimated by 16S rRNA gene quantities (Figure 3), which has the potential to realize greater substrate cleavage and consumption. This is supported by the reported phytase assay values prior to normalization (23.6 ± 13.2 nmol and 54.4 ± 36.5 nmol of P from the Ca-IP_6_ and the Ca-IP_6_+RE hotspots respectively). However, as the as the substrate availability to the enzyme is assumed to be the same in both samples, it is plausible that the populations from the two hotspots produced comparable amounts of phytate degrading enzymes per bacterium as seen in Figure 2. Finally, while the final product of phytases involve a lower order inositol phosphate (IP_5_-IP_1_) (Oh et al. 2001, Shin et al. 2001, Shim and Oh 2012), we detect no lower *myo*-inositol phosphates in our post-incubation HPLC results. A low IP_6_ availability will consequently result in small amounts of lower order *myo*-inositol phosphates which are prone to further dephosphorylation, as they are more easily accessible to the phytases and phosphatases present in the general population. Therefore, we hypothesize that the total functional capacity of the soil microbiota is able to clear the small amounts of lower order products quickly, resulting in concentrations of lower order *myo*-inositol phosphates below the detection limit of the HPLC system used. Overall, we take the combination of released Pi from Ca-phytate in the phytase assay and clearance of *myo*-inositol phosphate in the HPLC assay as an indication that the population from Ca-IP_6_-containing hotspots can degrade Ca-phytate at pH 7.2, albeit, slowly. The observations therefore suggest dephosphorylation of only the soluble fraction Ca-phytate present in an equilibrium with insoluble Ca-phytate, which appears unavailable to the enzyme.

The estimated bacterial abundance in each hotspot and bulk soil was measured via qPCR of the 16S rRNA gene. Results from the bulk soil are in agreement with other studies quantifying 16S rRNA in nutrient-deficient soils (Keshri et al. 2015), and as expected, significantly different bacterial numbers were recruited by the various hotspots (Figure 3). Total 16S rRNA numbers do not take bacterial turnover through death and degradation into account, nor predation of bacteria by other microorganisms (Rousk and Baath 2011). Furthermore, our data only estimates microorganisms from the bacterial kingdom containing the 16S rRNA gene, but especially in this sample supplied with both C and P, micro-eukaryotes (as fungi, protozoa, mites, nematodes) are likely also to be at play consuming a more palatable bacterial population (Jousset 2012, Otto et al. 2016). This may be the reason that lower numbers were recruited in the Ca-IP_6_+RE sample. The lowest number of bacteria were recruited to the Ca-phyate only treatment (Ca-IP_6_), which would suggest that this sample presents both challenging forms of P and C. Interestingly, it is also in this sample that the highest absolute copy numbers in *phoX* and *phoD* were observed, and which translated into high relative copy numbers (Figure 5). Finally, the high total number of absolute gene copies of phosphatase (*phoD* + *phoX* + *BPP*) in the RE treatment supports that the P-deficient soil used had pre-selected P-degrading functionality, since high potential to liberate P when C was provided was observed. The total population likely dilutes this, as the relative gene abundance shows a low copy number/16S rRNA ratio. The microcosm soil contains phytate, therefore small amounts of inherent precipitated phytate are likely to be present in the RE microcosm, helping also to explain the total bacterial abundance.

The organic P-cycling genes tested included *phoD* and *phoX* encoding alkaline phosphatases, and BPP, specific to the hydrolysis of Ca-phytate. *PhoD* is shown to exhibit phosphodiesterase ability and *phoX* is shown to act on phosphate monoesters (Roy et al. 1982, Gomez and Ingram 1995). Therefore, the presence of these genes represents an opportunity to cleave orthophosphate from lower order *myo*inositol phosphates. We hypothesized that the presence of organic P (Ca-phytate) would result in a recruitment and activity of microorganisms carrying organic P degradation-related genes. This response was, however, only seen in the Ca-IP_6_ hotspot, and only in absolute and relative abundance of *phoX* and *phoD*. Since both *phoD* and *phoX* abundance have been shown to respond to organic P and C, and the *phoD* abundance showed a particularly strong correlation to soil organic C (Ragot et al. 2016), these results are likely explained by the lack of easily accessible C and P in Ca-IP_6_. Looking specifically at the Ca-IP_6_ and the Ca-IP_6_+RE, the latter is supplied with a more diverse array of organic C and recruits sufficient microbial biomass to release orthophosphate from IP_6_ as measured in the non-normalized phytase assay (as reported) and the degradation of Ca-phytate as observed by HPLC. When orthophosphate becomes available via successful extracellular phosphatase activity, scavenger bacteria will also benefit and grow (*i.e*. those without alkaline phosphatase genes in their genome) and hence dilute the population carrying our genes of interest. Thus, in the CaIP_6_+RE hotspot where more total P is released, a lower relative gene copy per 16S rRNA gene would be expected as shown in Figure 5. While the relative abundance of the BPP being low in comparison to *phoD* and *phoX* has previously been shown (Neal et al. 2017, Neal and Glendining 2019), we did not see any difference in relative BPP abundance when compared between treatments (Figure 5). However, the detected phytase/phosphatase enzyme activity as well as the detected loss of IP_6_ in our HPLC assay support that alkaline phytase enzymes are active in our Ca-phytate-containing hotspots (Table 1, Figure 2). This challenges the view that BPP enzymes are main contributors to Ca-IP_6_ degradation in alkaline soil systems (Jorquera et al. 2013). There are, however, multiple explanations for this. First, the expression and downstream action of the BPP enzyme varies between treatments, which is supported by the BPP gene quantification showing equal relative gene copies in all populations (Figure 5), while the phytase assay detected orthophospahte-release only in Ca-IP_6_ and Ca-IP_6_+RE hotspot populations (Figure 2). Second, a C limitation drives the population evolution, necessitating only small amounts of the BPP enzyme to cleave the first phosphate off the inositol ring to render lower order *myo*-inositol phosphates theoretically available to other phosphatases (*i.e. PhoD* and *PhoX)* and *myo*-inositol dehydrogenases. Soil bacteria are able to utilize *myo*-inositol as a C source (Yoshida et al. 1997, Fry et al. 2001, Yebra et al. 2007), therefore, it is not unreasonable to assume that the recruited populations would utilize both the C and P from Ca-phytate. This would further support the lack of lower order *myo*-inositol phosphate products observed in our HPLC assay and suggests that the soils used are not only limited in bio-available P but also in bio-available C (Demoling et al. 2007). Third, the BPP primers employed do not cover enough sequence variability to capture the full BPP profile. Whilst there have been various attempts to develop degenerate BPP primers (Huang et al., 2009, Jorquera et al., 2013, Lim et al., 2007); a broad consensus remains however that the currently available primers are unable to capture the BPP profile believed to be present based on *in-vitro* evidence (Sanguin et al. 2016). Finally, there is the possibility that another novel or overlooked phytase acts in alkaline-buffered systems. Soil microorganisms in healthy, fertile soils provide a degree of functional redundancy (Jurburg and Salles 2015, Jia and Whalen 2020) however, the BPP is the only known microbial phytase to act under neutral or alkaline conditions. It is therefore not unreasonable to suggest that other enzymes are able to perform this task. Furthermore, as there is limited information available on the substrate specificity of the *PhoX* and *PhoD* in relation to phytate, and both have been shown to necessitate Ca^2+^ ions and function under alkaline conditions (Roy et al. 1982, Gomez and Ingram 1995), we speculate that these enzymes play a larger role in phytate degradation than previously shown. Further studies need to be undertaken to reveal the functional role of *PhoX* and *PhoD* in cycling of phytate in the soil environment.

To investigate the broad community differences between treatment populations in a time-sensitive manner, the catabolic profiles of the hotspot communities were subjected to BioLog^®^Eco plates. Here, 31 soil-specific substrates were monitored and provided initial insights into population differences, showing that the RE community was considerably different from the Ca-phytate-harboring communities (Ca-IP_6_ and Ca-IP_6_+RE), while the Ca-phytate-harboring communities in themselves showed a less distinct difference (Figure 6). These differences in community composition were further verified by 16S rRNA amplicon sequencing, where designation of OTU’s and SIMPER analysis provided deeper resolution. As such we showed that the hotspot bacterial communities were significantly different and less diverse than the soil bacterial community. A decrease in microbial community diversity is in agreement with previous studies on other hotspot environments, for example roots (Yousuf et al. 2012, Aislabie and Deslippe 2013) and fungal hyphae (Ghodsalavi et al. 2017, Hao et al. 2020) as compared to soil communities. As per SIMPER analysis, these differences were largely driven by the presence of Bacteriodetes at higher levels in RE hotspots receiving only root exudates, a high relative abundance of Firmicutes in Ca-IP_6_+RE hotspots, and relatively increased abundance of Actinobacteria in Ca-IP_6_ hotspots receiving only Ca-phytate (Figure 8A). Abundance of Actinobacteria in all hotspots of more than 50 % is in concordance with this phylum being dominant in bulk soils with limited organic matter and especially in soils abundant in phytate availability (Bell et al. 2014, Neal et al. 2017). In the present study the findings of Bacteroidetes in RE hotspots, is supported by Bacteroidetes being classified as copiotrophic groups commonly detected in C-enriched environments such as the rhizosphere (Yousuf et al. 2012, Aislabie and Deslippe 2013). Likewise, our findings of Firmicutes in the Ca-IP_6_+RE hotspots fit with previous studies showing that the phyla, and specifically the genus *Bacillus*, responds quickly to additions of organic material; (Fierer et al. 2007). Finally, our Ca-IP_6_ hotspot findings further compliment the findings by Sanguin et al. (2016) who employed a culture-independent 16S rRNA gene-DGGE study of the grass rhizosphere, and showed predominance of Proteobacteria and Actinobacteria in Na-phytate treated alkaline soils.

A majority of the phytate-hydrolyzing soil strains recognized presently have been isolated and/or identified using culture-dependent methods. These methods have largely yielded fast growing *Pseudomonas* and *Bacillus* strains from bulk soils (Richardson and Hadobas 1997, Idriss et al. 2002), and *Pseudomonas, Enterobacter, Pantoea*, and *Burkholderia* spp. from plant rhizospheres (Unno et al. 2005, Jorquera et al. 2008) When considering our OTU assignments, a specific increase of *Streptomyces* within the Actinobacteria phyla was evident. Furthermore, OTUs recognized as shared between Ca-IP_6_ and Ca-IP_6_+RE microcosms were Actinobacteria with the genera *Streptomyces, Promicromonospora* and *Nocardia* represented, indicating a genera of potential interest in the context of organic P cycling. A recent cloning and confirmation of a BPP in *Streptomyces* sp. UD42 (Boukhris et al. 2016) and a study of bacterioplankton (Valdespino-Castillo et al. 2017), as well as a search of the published full genome of *S. atratus* strain OK807 (UniProt), confirms that some *Streptomyces* contain phytases and phosphatases (including BPP and *phoD)* and emphasizes the potential of *Streptomyces* in phytate mineralization in an agricultural setting. Aside from putative phytase genes detected in selected *Streptomyces* species (Ghorbani-Nasrabadi et al. 2012), many of the OTU assignments we report (Table 2) have no literature supporting the presence of a known phytase enzyme, yet these OTU’s were shown to be abundant and specific in hotspots specifically containing Ca-phytate. Therefore, our findings highlights the current lack of knowledge surrounding the identity of phytate degraders in soil systems.

## 5. Concluding statements

The microcosm system developed in this paper was designed with the understanding that our ability to comprehensively dissect soil’s complex mechanisms and interactions remains limited. The intended use of this microcosm is to by-pass this lack of knowledge yet use the highly evolved soil systems’ inherent potential. Using the microcosm approach, we were able to show specific recruitment of taxonomically and functionally distinct bacterial populations using Ca-IP_6_ as a bait compound. Our results propose putting emphasis on Actinobacteria, and in particular, the genus *Streptomyces* when screening and testing for new biofertilizers for use to release orthophosphate from recalcitrant soil organic P. In addition, we propose that our microcosm platform could be extended to other substrates of interest. We further conclude that our microcosm design presents a new option for isolating microorganisms with potential to be effective as Ca-phytate mineralizers *in solum* and brings us a step closer to being able to isolate inherently competitive microorganisms with specific functional traits for targeted agricultural use.

## 6. Acknowledgments

A special thank you to Dorthe Ganzhorn for the technical support in the laboratory. This study was funded by innovation foundation Denmark, grant number 1308-00016B under the project Microbial biofertilizers for enhanced Crop availability of P pools in soil and waste (MiCroP).

