## Supplemental material for "A novel microcosm for recruiting inherently competitive biofertilizer-candidate microorganisms from soil environments"

Supplementary material

Sup. Table S1. primers used in this study

| Target genes | primers | Sequences (5’-3’) | references |
|---|---|---|---|
| *16S rRNA gene* | 341F | CCTAYGGGRBGCASCAG | (Yu, Lee, Kim, & Hwang, 2005) |
| | 806R | GGACTACNNGGGTATCTAAT | |
| *phoD* | ALPS-F730 | CAGTGGGACGACCACGAGGT | (Sakurai, Wasaki, Tomizawa, Shinano, & Osaki, 2008) |
| | ALPS-R1101 | GAGGCCGATCGGCATGTCG | |
| *phoX* | phoX2-F | GARGAGAACWTCCACGGYTA | (Valdespino-Castillo et al., 2014) |
| | phoX2-R | GATCTCGATGATRTGRCCRAAG | |
| *BPP* | BPP-F | GACGCAGCCGAYGAYCCNGCNITNTGG | (Huang et al., 2009) |
| | BPP-R | CAGGSCGCANRTCIACRTTRTT | |

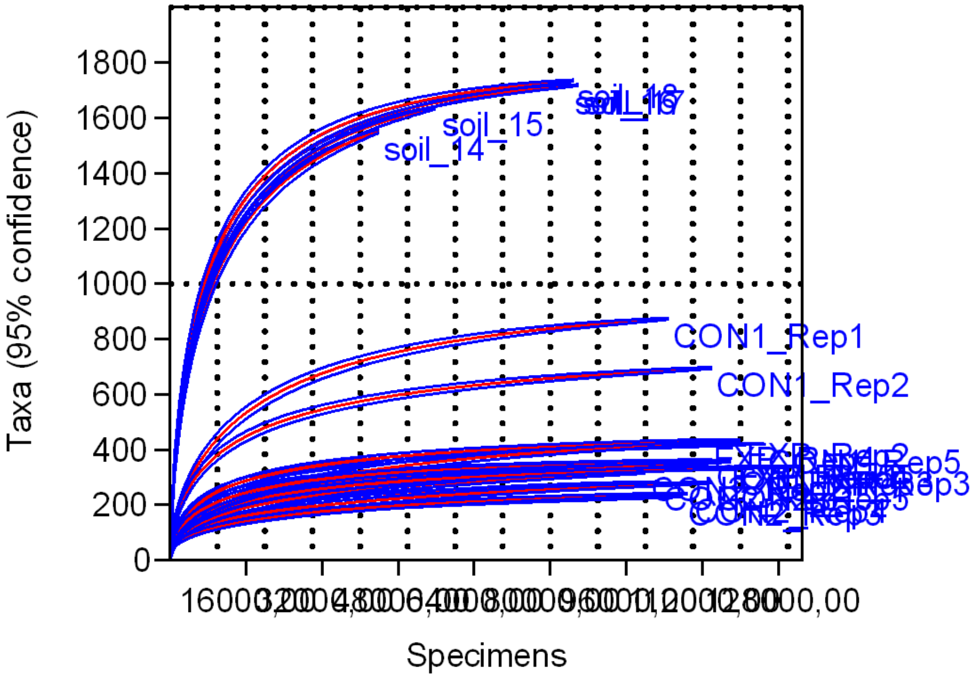


Sup. Figure S1. Rarefaction curve generated by the CLC program. Microcosms treatments are CON = Ca-phytate (Ca-$IP_6$), CON2 = root-exudate alone (RE), EXP = Ca- phytate combined with root-exudate (Ca-$IP_6$ + RE), and Soil = bulk soil. Five replicates per treatment were analyzed using 16S amplicon sequencing.

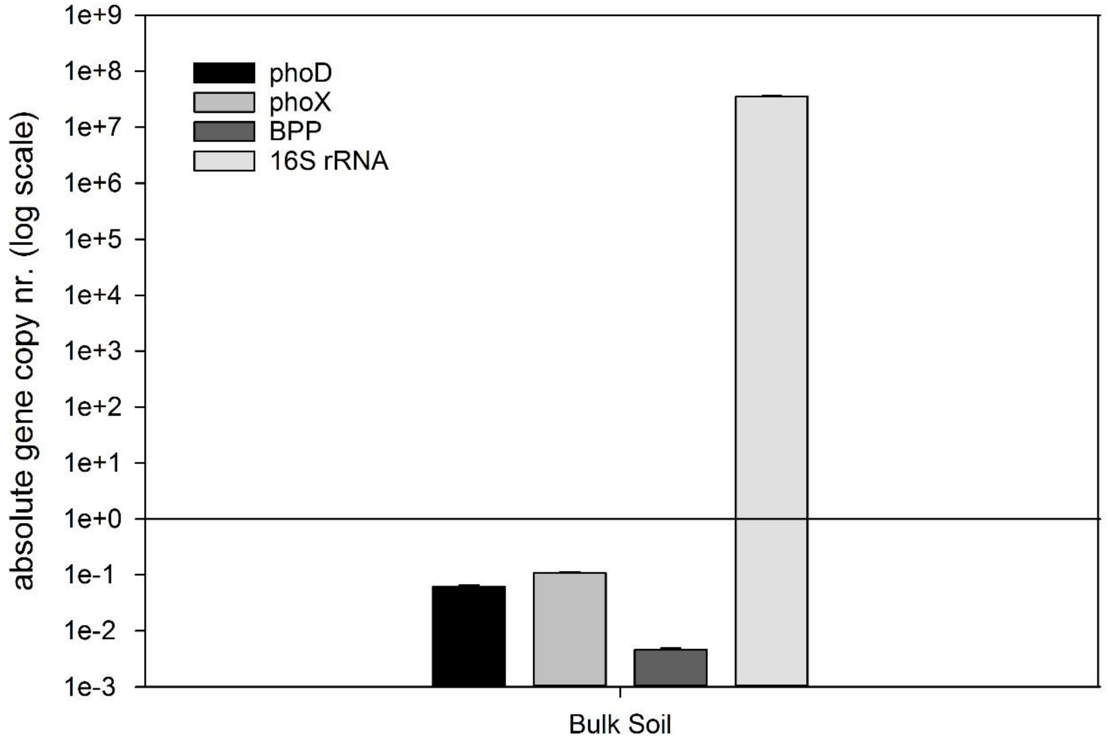


**Sup. Figure S2. Absolute abundance of marked genes found per gram bulk soil.** A. *phoX* = alkaline phosphatase, B. *phoD* = alkaline phosphatase, C. *BPP* = beta propeller phytase, *16S rRNA* = small ribosomal subunit gene. Error is shown in SEM.
